# Mammalian ATG9s drive the autophagosome formation by binding to LC3

**DOI:** 10.1101/2020.05.12.091637

**Authors:** Ting Zhang, Lixia Guo, Yanan Yang

**Affiliations:** Thoracic Disease Research Unit, Division of Pulmonary and Critical Care Medicine, Mayo Clinic Alix School of Medicine, Mayo Clinic, Rochester, Minnesota, 55905, USA; Department of Biochemistry and Molecular Biology; Mayo Clinic Alix School of Medicine, Mayo Clinic, Rochester, Minnesota, 55905, USA; Developmental Therapeutics and Cell Biology programs, Mayo Clinic Cancer Center, Mayo Clinic, Rochester, Minnesota, 55905, USA

## Abstract

The autophagosome is a membranous organelle that executes macroautophagy (call autophagy hereafter for the sake of brevity). Although the biogenesis of autophagosome depends on continuous acquirement of necessary membrane components to support its growth, the underlying mechanism has remained uncertain. Herein, we report that the mammalian ATG9 family members (maATG9s) play master roles in this process. We show that maATG9s directly localize to the growing autophagosomal membrane through their Ubiquitin-interacting motives (UIMs), which interact with the UIM docking site (UDS) on LC3, a component of the autophagosomal membrane. Interrupting the UIM-UDS interaction abrogates the maATG9s-LC3 binding, the localization of maATG9s to the growing autophagosome, and the autophagosome formation. Collectively, our findings show the first evidence that maATG9s act as primary autophagosomal membrane components to drive the autophagosome formation. As autophagy has been implicated in many human diseases, the identified maATG9s-LC3 interaction may serve as a target for manipulating pathological autophagosome formation.

## INTRODUCTION

The autophagosome is a double-membrane organelle that executes autophagy, a basic physiological process that transports certain cellular materials, such as damaged organelles, proteins, and lipids, to the lysosome for degradation and recycling, thereby maintaining the cellular material and energy balance^1–3^.

The autophagosome formation is a multi-stage process that depends on continuous acquirement of membrane components to support its growth^4–8^. In mammalian cells, a sub-organelle structure located at the end of the endoplasmic reticulum (ER), namely the omegasome, has been considered as the precursor of the autophagosome. Upon autophagic stimuli, the omegasome buds off the ER, localizes to the sites adjacent to the cellular contents that are destined for autophagic degradation, acquires necessary membrane, and grows into the phagophore (pre-autophagosome), which is morphologically an unclosed double-membrane structure under electron microscope. Next, the phagophore continues to acquire membrane components, expands, and eventually is closed to form a complete autophagosome. Then, the engulfed cellular contents can be transported to the lysosome for degradation, through a process involving the fusion of the autophagosome and the lysosome (to form the autolysosome).

How autophagosome continuously acquires its membrane remains poorly understood. In mammalian cells, two transmembrane maATG9s, including ATG9a and ATG9b, are thought to be responsible for this process by delivering membrane to the growing autophagosome. However, little mechanistic studies support such function for maATG9s. Notably, a previous report^9^ has shown that ATG9a neither stably associates with the autophagosome nor apparently localizes to the autophagosomal membrane, leaving it an open question whether maATG9s play significant roles in the autophagosomal membrane growth.

In the present study, we show unexpected findings that both members of maATG9s directly and extensively localize to the growing autophagosomal membrane through their interaction with LC3, a structural and functional component of the autophagosomal membrane^4, 5^. Mechanistically, we show that maATG9s contain previously unrecognized UIM motives, which mediate their binding to the UDS site on LC3. Consistent with these findings, interrupting the UIM-UDS interaction abrogates the maATG9s-LC3 binding, the localization of maATG9s to the growing autophagosome, and the autophagosome formation. These findings provide the first evidence that maATG9s are key components of the autophagosomal membrane, where they bind to LC3 to drive the autophagosome formation.

## RESULTS

### ATG9b co-localizes with endosome, Golgi, autophagosome and lysosome proteins

The maATG9 family consists of ATG9a and ATG9b. Previous studies have intensively focused on ATG9a but not ATG9b^10–18^. To understand the role for ATG9b, we expressed a FLAG-tagged ATG9b and performed confocal fluorescence microscopy studies, which show that ATG9b localizes to the endosome and the Golgi, as evidenced by its co-localization with early endosome antigen (EEA1) and the Golgi protein RCAS1^19^, respectively (Fig.1A-A’’’’ and B-B’’’’). These results are consistent with previously reported intracellular localization of ATG9a ^10–18^. On the contrary, ATG9b does not co-localize with the plasma membrane protein E-cadherin or the ER protein PDI^20, 21^ (Fig.1C-C’’’’ and D-D’’’’).

**Fig.1.**
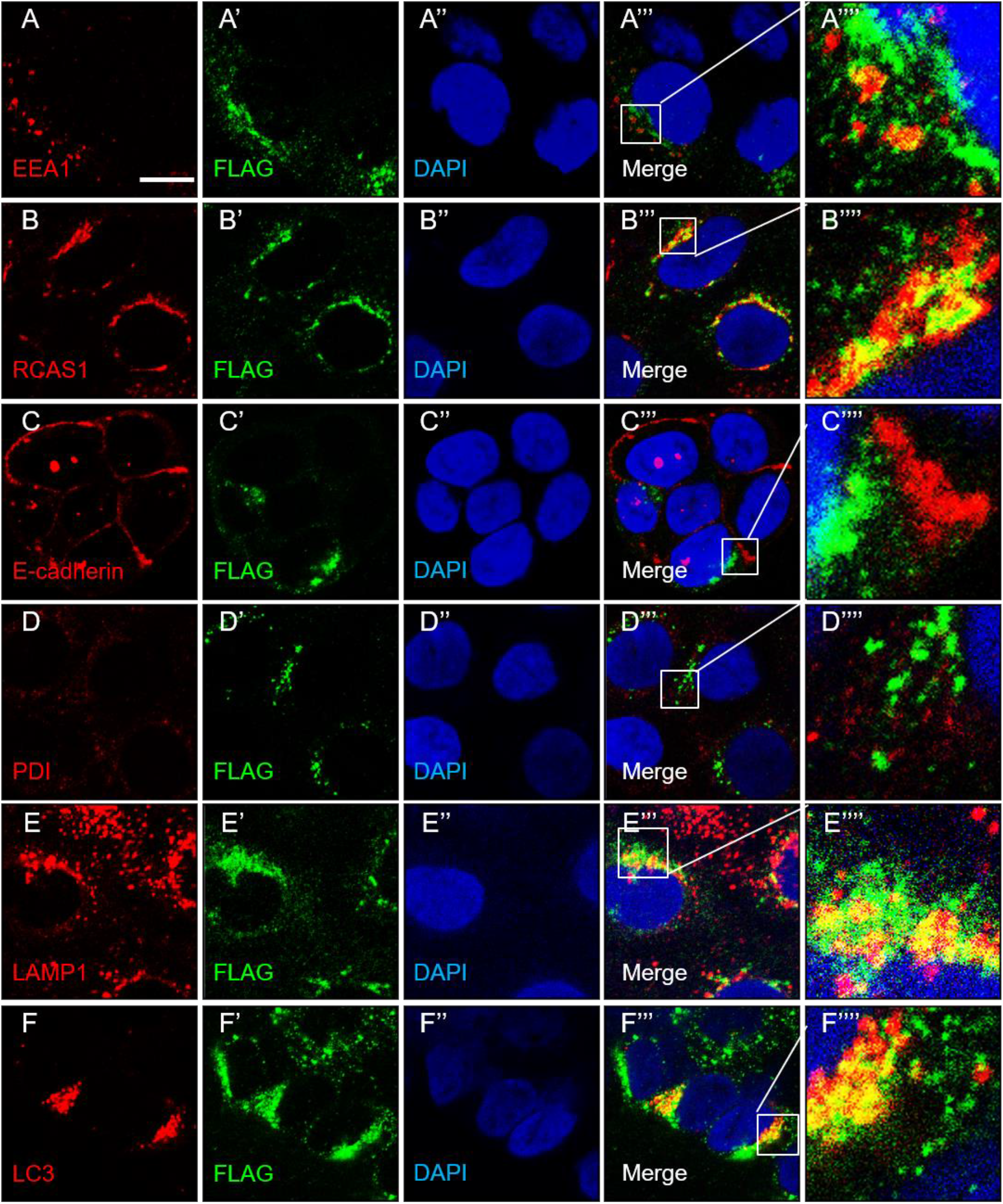
The intracellular localization of ATG9b. HCC827 human lung cancer cells stably expressing FLAG-tagged ATG9b (FLAG-ATG9b) were stained with antibodies against organelle markers, including EEA1 (A), RCAS1 (B), E-cadherin (C), PDI (D), LAMP1 (E), and LC3 (F), co-stained with anti-FLAG (A’, B’, C’, D’, E’, and F’) and DAPI (A’’, B’’, C’’, D’’, E’’, and F’’), and subjected to confocal fluorescence microscopy analysis. A’’’, B’’’, C’’’, D’’’, E’’’, and F’’’ show merged confocal fluorescence microscopy images. A’’’’, B’’’’, C’’’’, D’’’’, E’’’’, and F’’’’ shown blown-up images from white boxed areas from the merged images as indicated. Bar: 5 μm.

Notably, a large portion of ATG9b co-localizes with the lysosome protein LAMP1 (Fig.1E-E’’’’) and the autophagosome membrane protein LC3 (Fig.1F-F’’’’), suggesting that ATG9b may be a component of the autophagosome, which is distinct from the reported role for ATG9a^9^. To confirm this finding, we performed super-resolution structure illumination microscopy (SIM) studies. The results clearly show that a major part of the regenerated SIM structures contain co-localized ATG9b and LC3 (Supplementary Fig.1A and B; the white structures contain co-localized ATG9b and LC3).

To validate the above experiments that utilize exogenous FLAG-tagged ATG9b, we performed both confocal fluorescence microscopy (Supplementary Fig.1C-G) and SIM studies (Supplementary Fig.1H-K) for endogenous ATG9b and LC3. Consistent with our above findings, their extensive co-localization can be detected in all studies.

### The maATG9s co-localize with the autophagosome membrane proteins

Since previous studies have shown that ATG9a does not localize to the autophagosome membrane^9^, our above results suggest a possibility that ATG9a and ATG9b might differentially co-localize with LC3. If this is true, then it would suggest that ATG9b and ATG9a play distinct roles in the autophagosome formation. To address this possibility, we performed SIM studies for cells co-expressing RFP-tagged ATG9a (RFP-ATG9a) and GFP-tagged ATG9b (GFP-ATG9b) (Fig.2A-K). Strikingly, we found that both ATG9a and ATG9b similarly co-localize with LC3 (Fig.2E and F), suggesting that both of them localize to the autophagosome.

**Fig.2.**
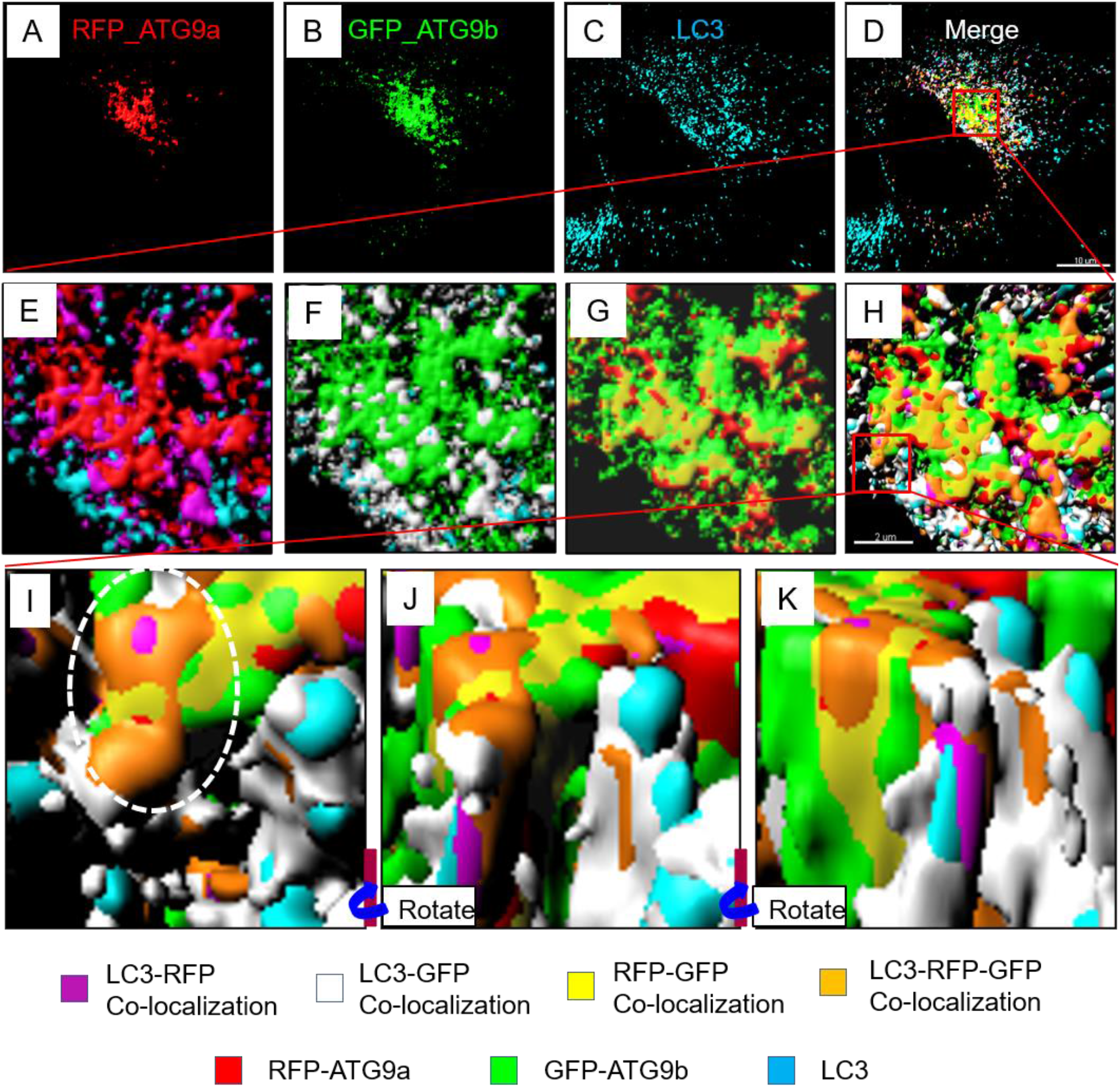
The co-localization of maATG9s with LC3 in SIM structures. H1299 human lung cancer cells were transiently co-transfected with RFP-ATG9a and GFP-ATG9b, stained with anti-LC3, and subjected to SIM structure analysis. A-C: regenerated single channel surfaces for RFP (A), GFP (B), and LC3 (C), respectively. D: the merged regenerated SIM structures, including the three single channel surfaces shown in A-C, the co-localiztion of any two of the three surfaces, and the co-localization of all three surfaces together. Bar: 10 μm. E-H: the blown-up for a red boxed area from D as indicated. H include all regenerated SIM structures, including the three single channel surfaces shown in A-C, the co-localiztion of any two of the three surfaces, and the co-localization of all three surfaces together. E-G show any two of the three single channel surfaces and their co-localization. Bar: 2 μm. I: the blown-up for a red boxed area from H as indicated. The white dash line circled area in I shows a representative structure (brown) that is triple positive for RFP, GFP, and LC3. J and K: the same structures in I were rotated to provide more comprehensive views.

Furthermore, in depth analysis of the SIM results reveals that ATG9a, ATG9b, and LC3 can co-localize to the same structure (Fig.2H-K, the white dash line-circled area in Fig.2I shows a representative triple positive brown-colored structure), suggesting that the co-localizations of ATG9a or ATG9b with LC3 are not mutually exclusive. Therefore, ATG9a and ATG9b may utilize similar mechanism(s) and act in a cooperative manner to co-localize with LC3, and it is possible that they exert similar functions in the autophagosome formation.

It should be noted that, although LC3 is frequently used as an autophagosome membrane marker, it does have non-autophagosome localization^4, 5^. As such, the above-shown maATG9s-LC3 co-localization may not necessarily indicate autophagosome localization of the maATG9s. To address this point, we performed SIM analysis (Fig.3A-K) for RFP-ATG9a, GFP-ATG9b, and WIPI2, which is a phagophore membrane protein^4, 5^. Similar to our above-shown observations, ATG9a and ATG9b, individually or together, can co-localize with WIPI2 (the white dash line-circled areas in Fig.3I-K show representative co-localizations).

**Fig.3.**
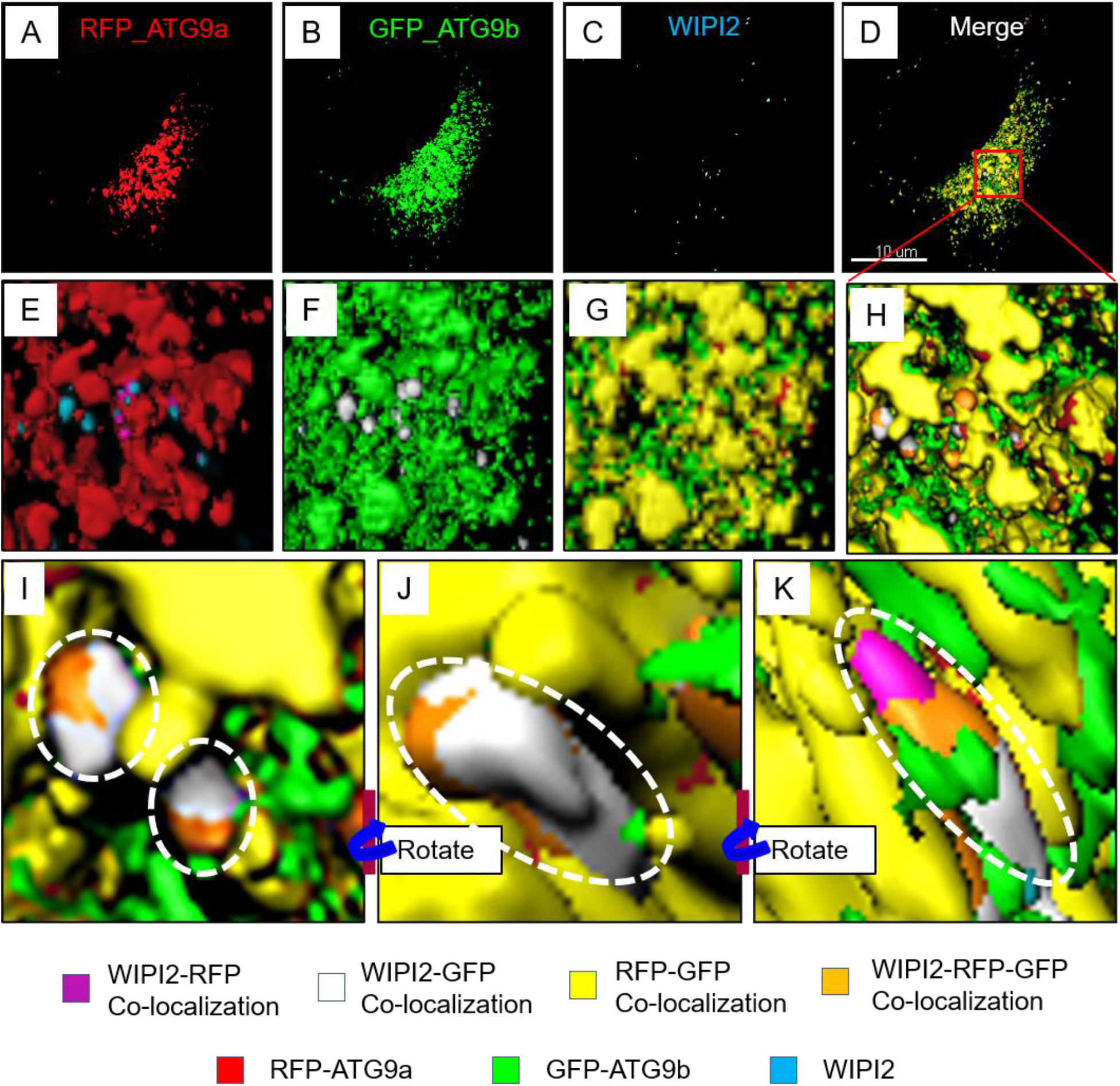
The co-localization of maATG9s with WIPI2 in SIM structures. H1299 human lung cancer cells were transiently co-transfected with RFP-ATG9a and GFP-ATG9b, stained with anti-WIPI2, and subjected to SIM structure analysis. A-C: regenerated single channel surfaces for RFP (A), GFP (B), and WIPI2 (C), respectively. D: the merged regenerated SIM structures, including the three single channel surfaces shown in A-C, the co-localiztion of any two of the three surfaces, and the co-localization of all three surfaces together. Bar: 10 μm. E-H: the blown-up for a red boxed area from D as indicated. H include all regenerated SIM structures, including the three single channel surfaces shown in A-C, the co-localiztion of any two of the three surfaces, and the co-localization of all three surfaces together. E-G show any two of the three single channel surfaces and their co-localization. I: the blown-up for a red boxed area from H as indicated. The white dash line circled areas in I show representative structures (brown) that are triple positive for RFP, GFP, and WIPI2. J and K: the same structures in I were rotated to provide more comprehensive views.

Intriguingly, we note that there is almost no SIM structures only positive for WIPI2 (Fig.3H-K), but there are abundant SIM structures that are only positive for LC3 (blue structures Fig.2H-K). Because LC3 is present on the membrane of both phagophore and autophagosome, whereas WIPI2 is only present on phagophore membrane, this finding suggests that ATG9a and ATG9b start entering the autophagosome membrane at the phagophore stage.

### The maATG9s are components of the mammalian autophagosomal membrane

To more precisely understand the localization of maATG9s to the autophagosome, we labeled them with an APEX2 gene, which encodes for a peroxidase and has been used as a molecular tag for visualizing intracellular localization of interested proteins under electron microscope^22–24^. Our results show that the APEX2-labeled ATG9a and ATG9b exhibit apparent and selective localization to the membrane of both phagophore (Fig.4A-D) and autophagosome (Fig.4E-H). These findings are consistent with our above-discussed SIM results.

**Fig.4.**
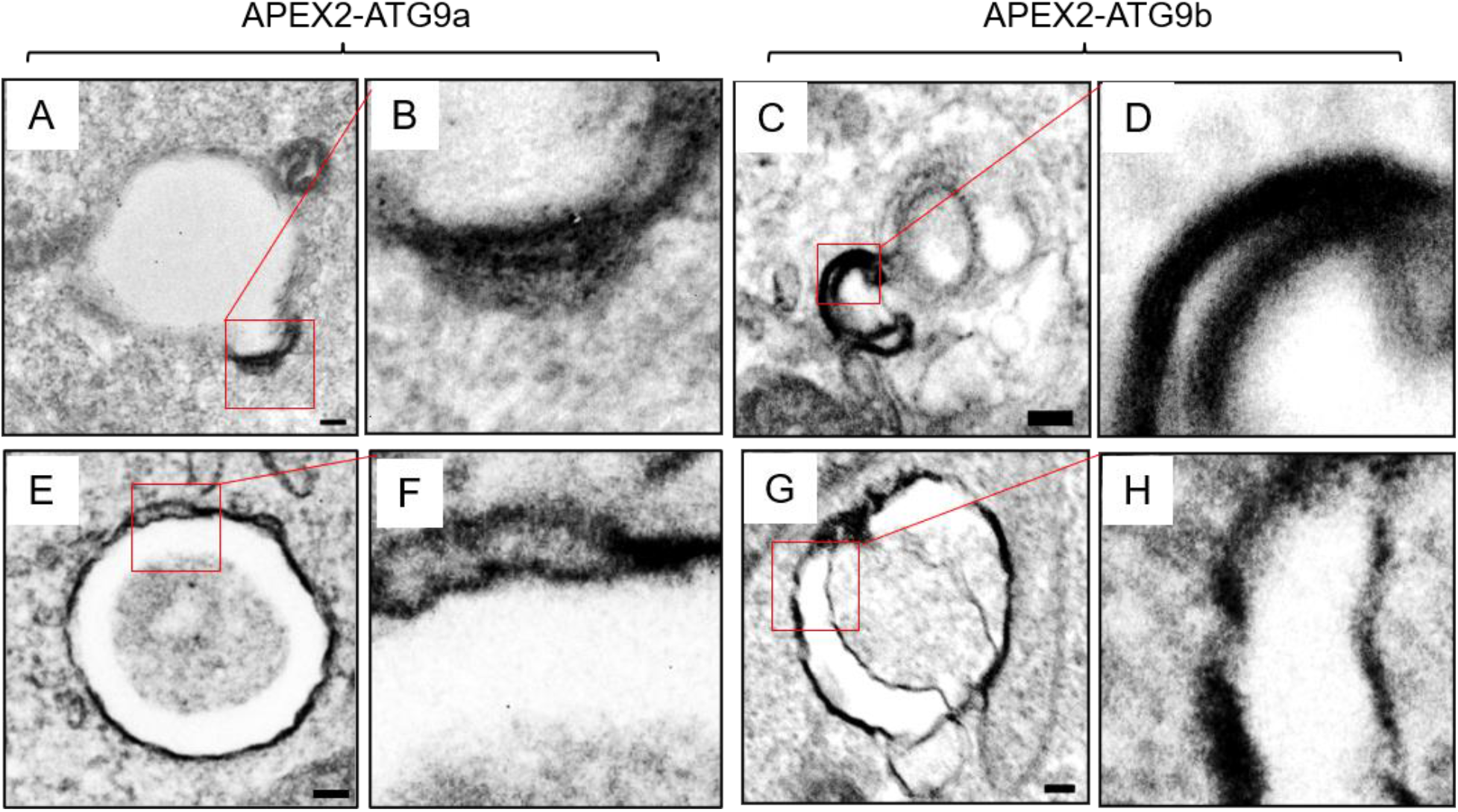
The autophagosome membrane localization of msATG9s. H1299 cells were transiently transfected with APEX2-ATG9a or APEX2-ATG9b and subjected to transmission electron microscopy analysis. The dark black areas are those exibiting APEX2 enzymatic activities, indicating the presence of APEX2-ATG9a or APEX2-ATG9b. A-D: representative images for phagophores in cells expressing APEX2-ATG9a (A and B) or APEX2-ATG9b (C and D). B and D are blown-ups of the red boxed areas from A and C, respectively. Bars: 100 nm. E-H: representative images for autophagosomes in cells expressing APEX2-ATG9a (E and F) or APEX2-ATG9b (G and H). F and H are blown-ups of the red boxed areas from E and G, respectively. Bars: 100 nm.

Next, by utilizing a serial block face scanning electron microscopy (SBF-SEM), we were able to more comprehensively understand the localization of the APEX2-labeled maATG9s to the autophagosomal membrane of a single autophagosome in a 3-dimensional (3D) setting (Fig.5A-O). Regenerated 3D models from the SBF-SEM results show that the majority of the autophagosomal membrane contain the APEX2-labeled ATG9a (Fig.5P-R) and ATG9b (Supplementary Fig.2A-C). Interestingly, neither of them covers 100% of the autophagosome, again suggesting that their autophagosome membrane localizations are not mutually exclusive. Collectively, our above results demonstrate that the maATG9s are components of the autophagosome membrane.

**Fig.5.**
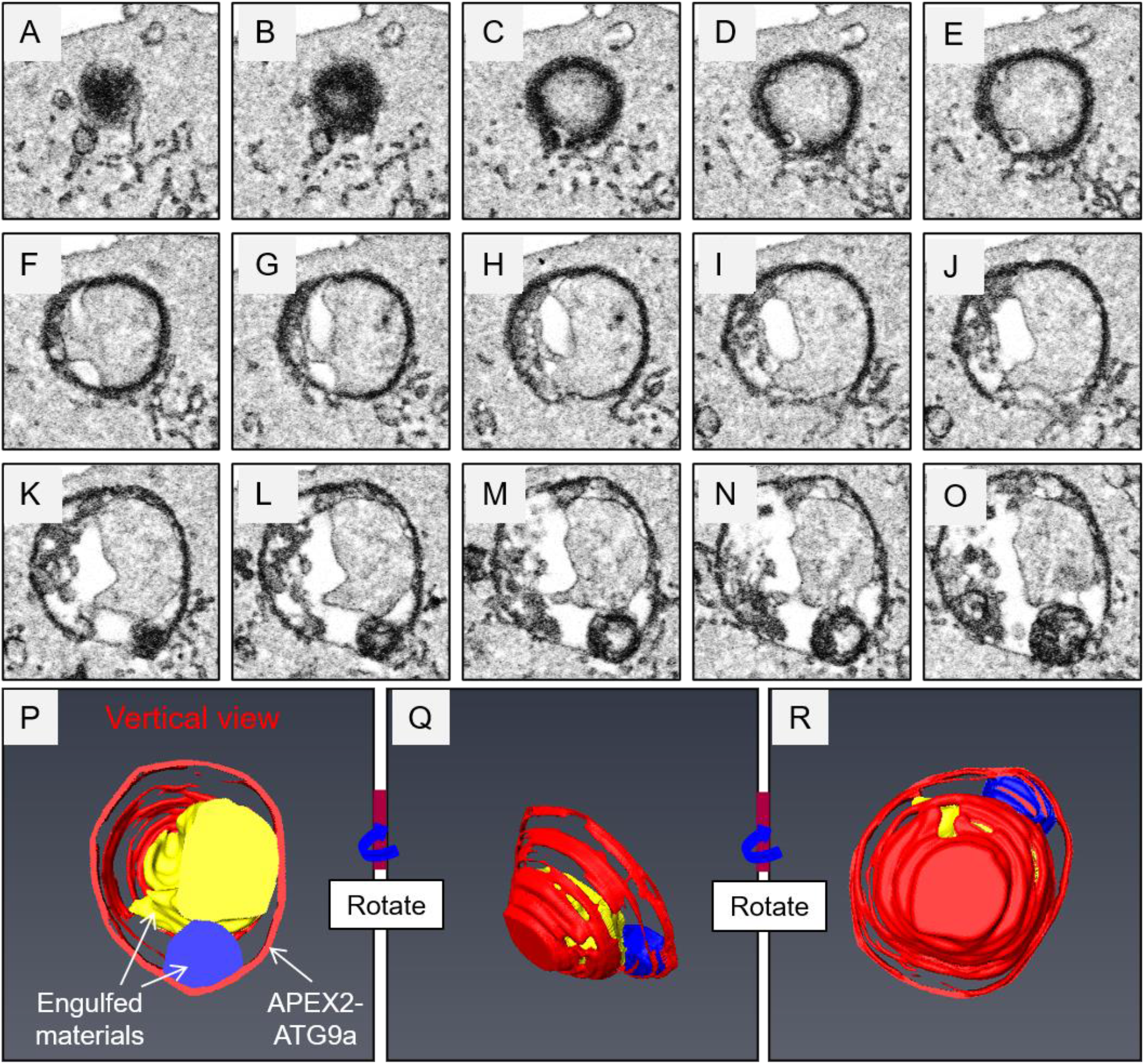
The autophagosome membrane localization of ATG9a in the SBF-SEM structure. H1299 cells were transiently transfected with APEX2-ATG9a and subjected to SBF-SEM analysis. The dark black areas are those exibiting APEX2 enzymatic activities, indicating the presence of APEX2-ATG9a. A-O: representative serial SBF-SEM images for an autophagosome. Bar: 100 nm. P: the regenerated SBF-SEM autophagosome structure using images shown in A-O. Q and R: the same structures in P were rotated to provide more comprehensive views.

### The maATG9s directly bind to LC3

Because the maATG9s co-localize with LC3, we suspect that they may directly interact with each other. In support of this idea, proximity ligation assays, which have been used to investigate in situ protein-protein binding^25, 26^, show abundant maATG9s-LC3 binding within the cells (Supplementary Fig.3A-D). Furthermore, our immunoprecipitation (IP)-Western blotting experiments show that both ATG9a and ATG9b co-immunoprecipitate with LC3 (Fig.6A and B).

**Fig.6.**
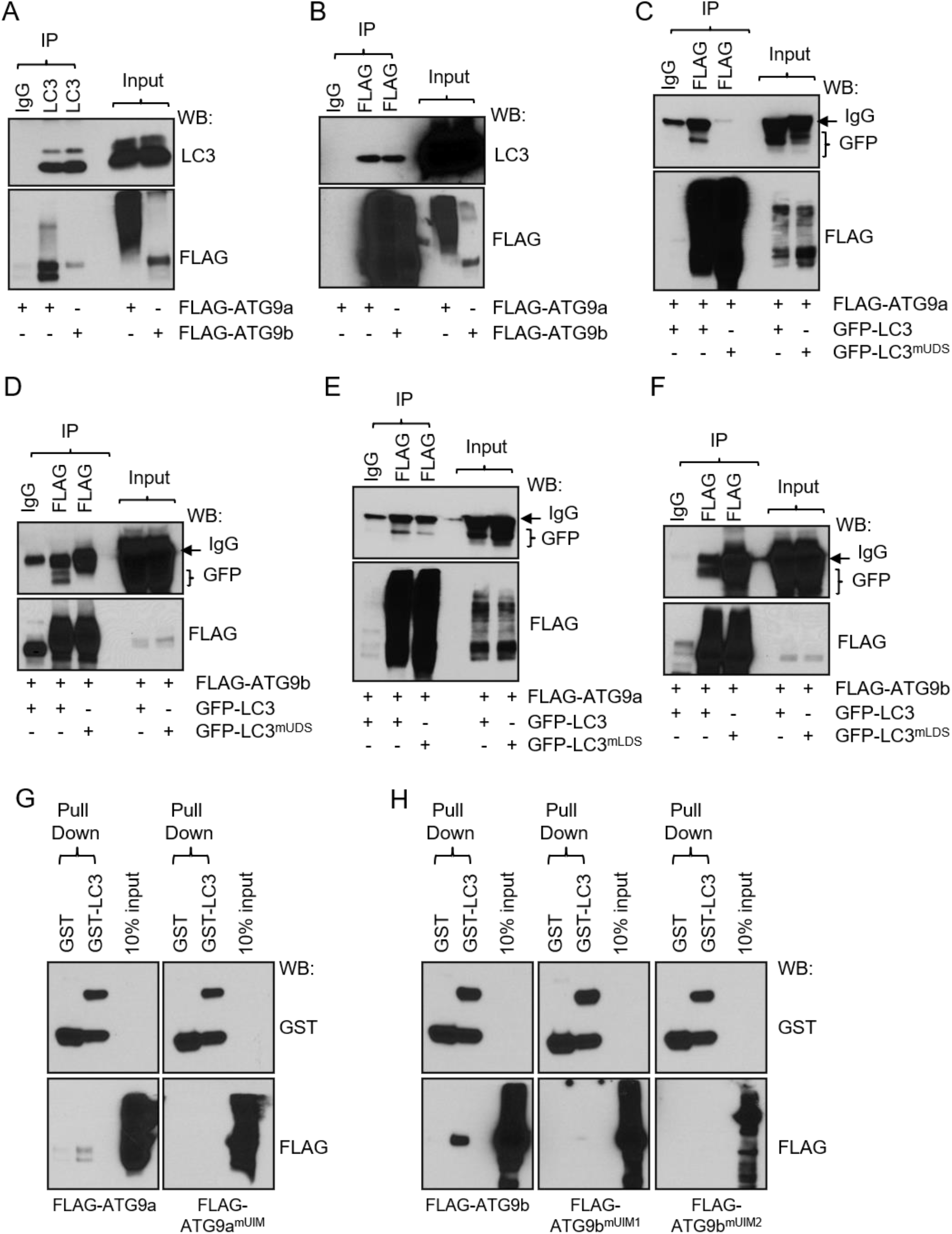
The UIMs of maATG9s bind to the UDS site on LC3. A and B: IP-Western blots for H1299 cells transfected with FLAG-ATG9a or ATG9b as indicated. Input: 10%. C and D: IP-Western blots for H1299 cells transfected with FLAG-ATG9a or ATG9b, and co-transfected with GFP-LC3 or GFP-LC3^mUDS^ (the UDS-depleted LC3), as indicated. Input: 5%. E and F: IP-Western blots for H1299 cells transfected with FLAG-ATG9a or ATG9b, and co-transfected with GFP-LC3 or GFP-LC3^mLDS^ (the LDS-depleted LC3), as indicated. Input: 5%. G and H: GST pull down assays using purified GST-LC3, FLAG-ATG9a, FLAG-ATG9a^mUIM^, FLAG-9b, FLAG-ATG9a^mUIM1^, and FLAG-ATG9a^mUIM2^ as indicated. Input: 10%.

LC3 has been shown to bind to other proteins through its LDS (LC3-interacting motif docking site) or UDS (Ubiquitin-interacting motif docking site) sites^4, 5, 27^. Interestingly, inactivation mutation^4, 5, 27^ of the UDS site (Fig.6C and D), but not the LDS site (Fig.6E and F), abrogates the binding between maATG9s and LC3.

Protein sequence analysis reveal that ATG9a has a ubiquitin-interacting motif (UIM), and ATG9b has two UIMs (Supplementary Fig.4A and B). These UIMs are conservative across species, but their locations on the two maATG9s are distinct. To determine whether these UIMs mediate the binding of maATG9s to LC3, we performed mutagenesis experiments to deplete each of them. Unfortunately, depletion of the UIMs dramatically destabilized ATG9b protein (Supplementary Fig.5A and B) and did not allow us to conclude their necessity. To address this problem, we performed GST-pull down experiments, which show that depletion each of the UIMs from ATG9a and ATG9b abrogates their binding to LC3 (Fig.6G and H), demonstrating that these UIMs are essential for maATG9s-LC3 binding, presumably through their direct interactions with the UDS site on LC3.

### The maATG9s localize to the autophagosome membrane to drive the autophagosome formation

On the basis of our above findings, we determined the functional role for the maATG9s-LC3 interaction in the autophagosome formation. By performing SIM studies, we found that depletion of the UIMs of ATG9a and ATG9b almost completely abrogates their co-localization with WIPI2 (Fig.7) or LC3 (Supplementary Fig.6), demonstrating that the maATG9s-LC3 binding is essential for maATG9s to localize to the autophagosome membrane.

**Fig.7.**
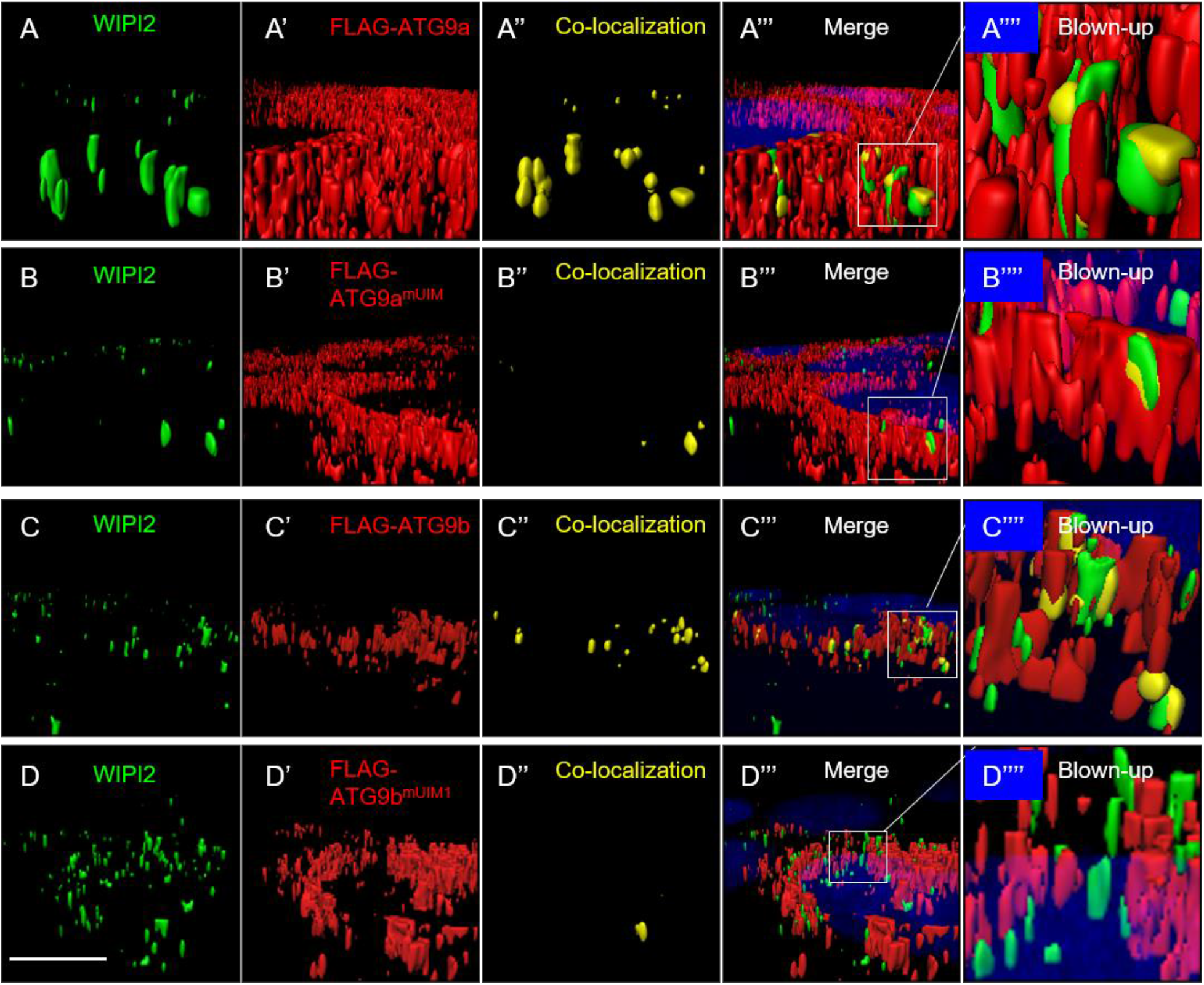
The localization of maATG9s to the phagophore depends on UIM motives. H1299 human lung cancer cells were transiently transfected with FLAG-ATG9a (A-A’’’’), FLAG-ATG9a^mUIM^ (B-B’’’’), FLAG-ATG9b (C-C’’’’), and FLAG-ATG9b^mUIM1^ (D-D’’’’), stained with WIPI2 (A, B, C, and D) and FLAG (A’, B’, C’, and D’), and subjected to SIM structure analysis. A, B, C, D, A’, B’, C’, and D’: regenerated single channel surfaces for WIPI2 and FLAG as indicated. A’’, B’’, C’’, and D’’: regenerated surfaces for the co-colocalizations of WIPI2 and FLAG. A’’’, B’’’, C’’’, and D’’’: the merged SIM structures. DAPI staining was included to show the nuclei. A’’’’, B’’’’, C’’’’, and D’’’’: the blown-up for the white boxed areas from A’’’, B’’’, C’’’, and D’’’ as indicated. Bar: 10 μm.

Next, we knocked out ATG9a gene by utilizing a CRISPR-cas9 approach, and re-expressed ATG9a or the UIM-depleted ATG9a (ATG9a^mUIM^) in the ATG9a knockout cells (Fig.8A). By quantitating the numbers of phagophore, autophagosome, and autolysosomes (collectively known as autophagic vacuoles or “AVs”), we found that knockout of ATG9a significantly deplete the AVs, and re-expressing ATG9a but not ATG9a^mUIM^ restore the number of AVs (Fig.8B and C), indicating that the binding of ATG9a to LC3 is required for the autophagosome formation.

**Fig.8.**
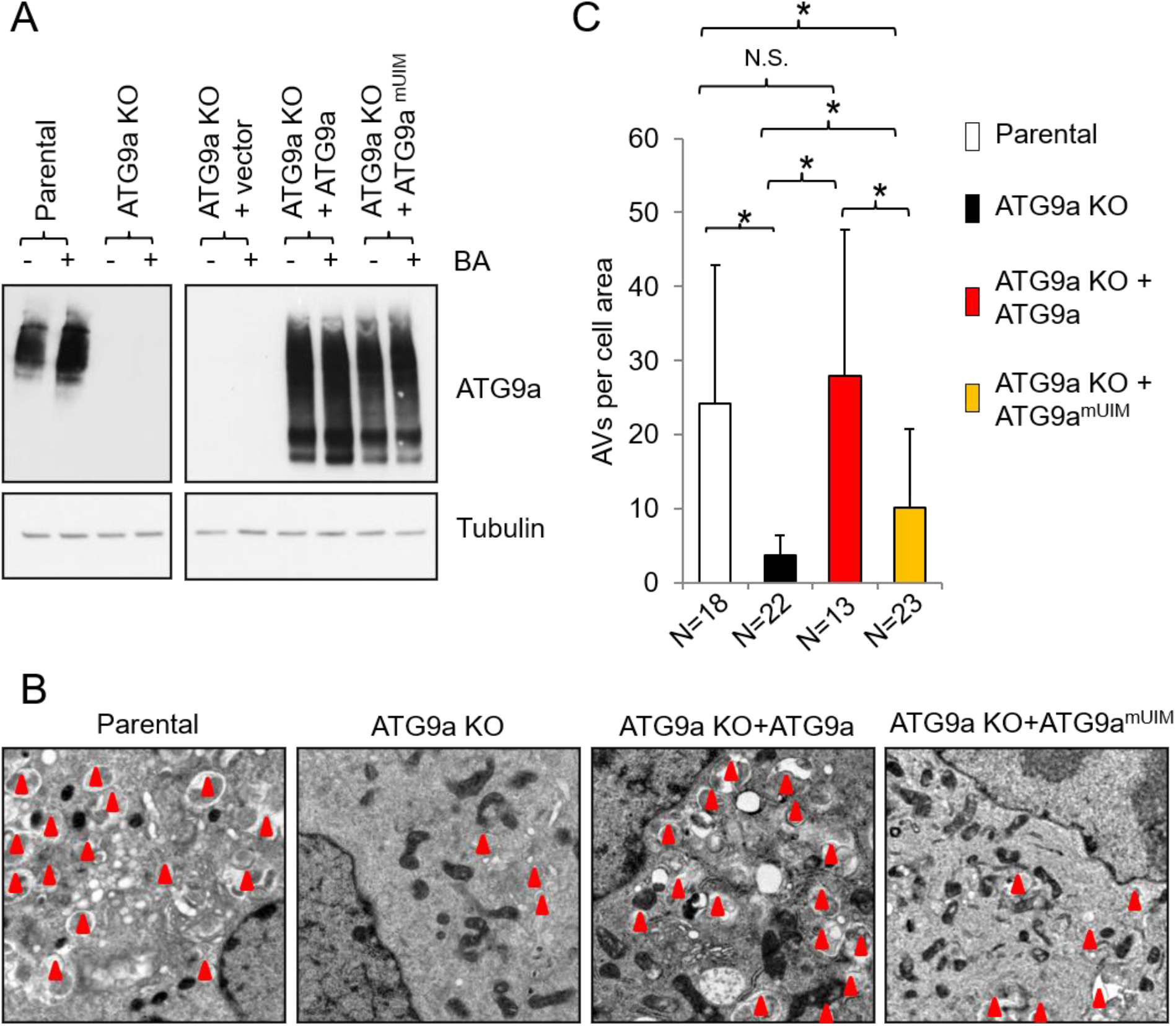
ATG9a drives the autophagosome formation through its UIM motif. A: Western blots for ATG9a knockout H1299 cells (ATG9a KO) and the ATG9a KO cells transiently transfected with vector, ATG9a, or ATG9a^mUIM^ in the presence or absence of bafilomycin A1 (BA, 500 nM, treated for 6 hours) as indicated. B: representative electron microscopy images for parental and ATG9a KO H1299 cells, and ATG9a KO cells transfected with ATG9a or ATG9a^mUIM^ in the presence of BA (BA, 500 nM, treated for 6 hours) as indicated. Red triangles indicate AVs. C: quantitation of the AVs in parental and ATG9a KO H1299 cells, and ATG9a KO cells transfected with ATG9a or ATG9a^mUIM^ in the presence of BA (BA, 500 nM, treated for 6 hours) as indicates. * indicates t-test p<0.05. N.S. indicates t-test p>0.05 (not statistically significant).

## DISCUSSION

In the present study, our results collectively reveal previously unrecognized roles for maATG9s as important components of the mammalian autophagosome membrane, where they act as drivers of the autophagosome formation. Our results are quite different from a previous report that ATG9a does not localize to the autophagosome membrane^9^. The reason for such discrepancy is unclear. Notably, that report was based on the results from immune electron microscopy studies, which largely depends on the quality of the antibody. Therefore, it is possible that the antibody used in that report was not sensitive enough to detect the autophagosome membrane-associated ATG9a. In addition, the techniques we used are quite different from those used in that report, such as the SIM, APEX2-labeling, and SBF-SEM techniques, (Figs.2-5, and Supplementary Figs.1 and 2), which have emerged in recent years as novel cell biology tools for visualizing intracellular protein localizations and functions. These advanced techniques may also contribute to our successful detection of the localization of maATG9s to the autophagosome membrane.

More importantly, our results identify a key interaction between maATG9s and LC3 (Fig.6), which plays essential roles in the localization of maATG9s to the autophagosome membrane and their ability to promote the autophagosome formation. We show that interrupting this interaction abrogates the autophagosome formation (Figs.7 and 8, and Supplementary Fig.6), suggesting that it may be useful within certain physiological or pathological contexts. For instance, misregulation of the autophagosome formation and autophagy has been widely implicated in human diseases, ranging from developmental defects to aging-associated diseases^1–3^, such as cancer, fibrosis, Alzheimer’s disease, Parkinson’s disease, Vici syndrome, and etc‥ Therefore, the maATG9s-LC3 interaction may serve as a novel target for developing therapeutic approach for manipulating disease-associated autophagosome formation.

It should be noted that, ATG9 has also been studied in other species, for example the yeast. Unlike, mammalian cells, yeast only has a single ATG9 gene. It forms a complex with other ATGs on the membrane of mitochondria to promote the formation of the PAS site, which is roughly equivalent to the phagophore in mammalian cells^4, 5^. Similar to our findings, the yeast ATG9 localizes to the autophagosome membrane^28^, but the underlying mechanism remains incompletely understood. It would be interesting for future studies to determine whether the ATG9-LC3 interaction similarly promotes the localization of yeast ATG9 to the autophagosome membrane.

## MATERIALS AND METHODS

### Cell lines, cell culture, and transfection

HCC827 and H1299 cells were gifts from Jonathan Kurie MD (The University of Texas MD Anderson Cancer Center). All cells were cultured in RPMI-1640 medium (purchased from Mediatech; catalog number MT10-040-CVRF) supplemented with 10% fetal bovine serum (purchased from Gibco; catalog number 26140079) in the absence of antibiotics. Transfection was performed by using Lipofectamine 2000 transfection reagent (purchased from Life Technologies; catalog number l3000015) and following the protocol provided by the manufacturer.

### DNA plasmid constructs and mutagenesis experiment

ATG9b cDNA was purchased from Origene (catalog number MR217776) and was sub-cloned into pCMV10-3xFLAG and pEGFP vectors. APEX2 cDNA was a gift from Mark McNivin Ph.D. (Mayo Clinic) and was subcloned into the pCMV10 vector. pMXs-puro-RFP-ATG9a plasmid (purchased from Addgene, catalog number 60609), from which Atg9a cDNA was amplified and subcloned into pCMV10-3xFLAG vector. pEGFP-LC3 (human) plasmid construct was purchased from Addgene (catalog number 24920). All mutant ATG9a, ATG9b, and LC3 plasmid constructs used in our studies were generated by performing mutagenesis experiments, using the Q5 quick mutagenesis kit (purchased from New England BioLabs, catalog number E0554s) and following the protocol provided by the manufacturer. All mutagenic primers are described in Supplementary Table 1.

### Antibodies

Information for the antibodies used in this study is listed in Supplementary Table 2.

### Chemicals and reagents

ProLong® Gold Antifade Mountant with DAPI was purchased from Invitrogen (P36931). Bafilomycin A1 (BA) was purchased from Cell Signaling Technologies (catalog number 54645). Purified GST peptide was purchased from Genscript (catalog number 202039). Purified GST-tagged LC3 protein was purchased from Ubiquigent (catalog number 60-0111-500). All other chemicals and reagents were purchased from Sigma unless specified.

### Proximity ligation assay (PLA assay)

The PLA assay was performed to determine the in situ binding between LC3 and ATG9a or ATG9b by using the Duolink® In Situ Red Starter Kit purchased from Sigma Aldrich (catalog number DUO92101), following the protocol provided by the manufacturer.

### GST pull-down assay

The GST pull down assay was performed by using the Pierce™ GST Protein Interaction Pull-Down Kit (catalog number 21516), following the protocol provided by the manufacturer. Briefly, the FLAG-tagged ATG9a, ATG9b, and their mutants were transiently transfected into H1299 cells. Protein lysates were prepared, and 5 mg protein lysates from each transfectant was incubated with 5 μg purified GST peptide or GST-tagged LC3 protein at 4 degree for overnight.

### Statistics

Statistical significance was determined by student’s *t* tests using the Microsoft excel software. When p values are less than 0.05, the results are considered statistically significant.

### Microscopy

All microscopy studies (confocal fluorescence microscopy, super resolution structure illumination microscopy, and electron microscopy) were performed at the Mayo Microscopy and Cell Analysis Core. The microscopes used in our studies include LSM-780 inverted confocal microscope (with Zeiss’s Zen software), Elyra PS.1 Super Resolution (Carl Zeiss Microscopy, LLC), 1400 TEM Transmission Electron Microscope (JEOL), and a volume scope scanning electron microscope (FEI). Super resolution structure illumination analysis was performed by using the Imaris 8 software (Oxinst), and structure regeneration from the SBF-SEM results was performed by using the Amira 2019 software (FEI).

### Preparation of whole cell protein lysate, immunoprecipitation, and Western blotting

To prepare whole cell protein lysate, cells were briefly washed with ice cold PBS (purchased from Mediatech; catalog number MT21-040-CVRF) and then directly lysed in RIPA buffer supplemented with proteinase inhibitor cocktail, PMSF, and sodium orthovanadate (all were from a RIPA lysis buffer system purchased from Santa Cruz; catalog number sc-24948A) following the protocol provided by the manufacturer. Immunoprecipitation for the protein lysate was performed by using SureBeads™ Protein G or A Magnetic Beads (purchased from Bio-Rad; catalog numbers1614023 and 1614013). Western blotting experiments were performed by using precast TGX 4-20% gels (purchased from Bio-Rad; catalog number 4561096) and the Trans-Blot® Turbo™ Rapid Transfer System (purchased from Bio-Rad; catalog number 1704150). Protein bands were visualized by incubating with ECL substrates (purchased from Bio-Rad; catalog number 1705061). The uncut gels for Western blots are shown in Supplementary Figs.7 and 8.

## ACKNOWLEDGEMENTS

This work was partly supported by the Mayo NIH relief award (CA218109A1relief), the Mayo Center for Biomedical Discovery cancer and aging platform award (93059043), and Mayo Cancer Center Developmental Therapeutics program pilot awards to Y.Y. pEGFP-LC3 (human) was a gift from Toren Finkel (Addgene plasmid # 24920; http://n2t.net/addgene:24920; RRID:Addgene_24920; reference 29). pMXs-puro-RFP-ATG9A was a gift from Noboru Mizushima (Addgene plasmid # 60609; http://n2t.net/addgene:60609; RRID:Addgene_60609; reference 30). The Mayo Microscopy and Cell Analysis Core was utilized for some of the experiments performed in this report, and we are grateful for the technical assistance from the core facility staff members, including Trace Christensen, Bingquan Huang, Kyle Howell, Jon Charlesworth, Duha Vang, Scott Gamb, and Duane Deal. We thank Jonathan Kurie MD (The University of Texas MD Anderson Cancer Center) and Edward Leof PhD (Mayo Clinic) for providing cell lines as gifts to us.

## AUTHOR CONTRIBUTIONS

Y.Y. conceived the project. L.G., T.Z., and Y.Y. performed experiments and collected and analyzed data. T.Z., L.G., and Y.Y. wrote and approve the paper.

## SUPPLEMENTARY FIGURE LEGENDS

**Supplementary Fig. 1.**
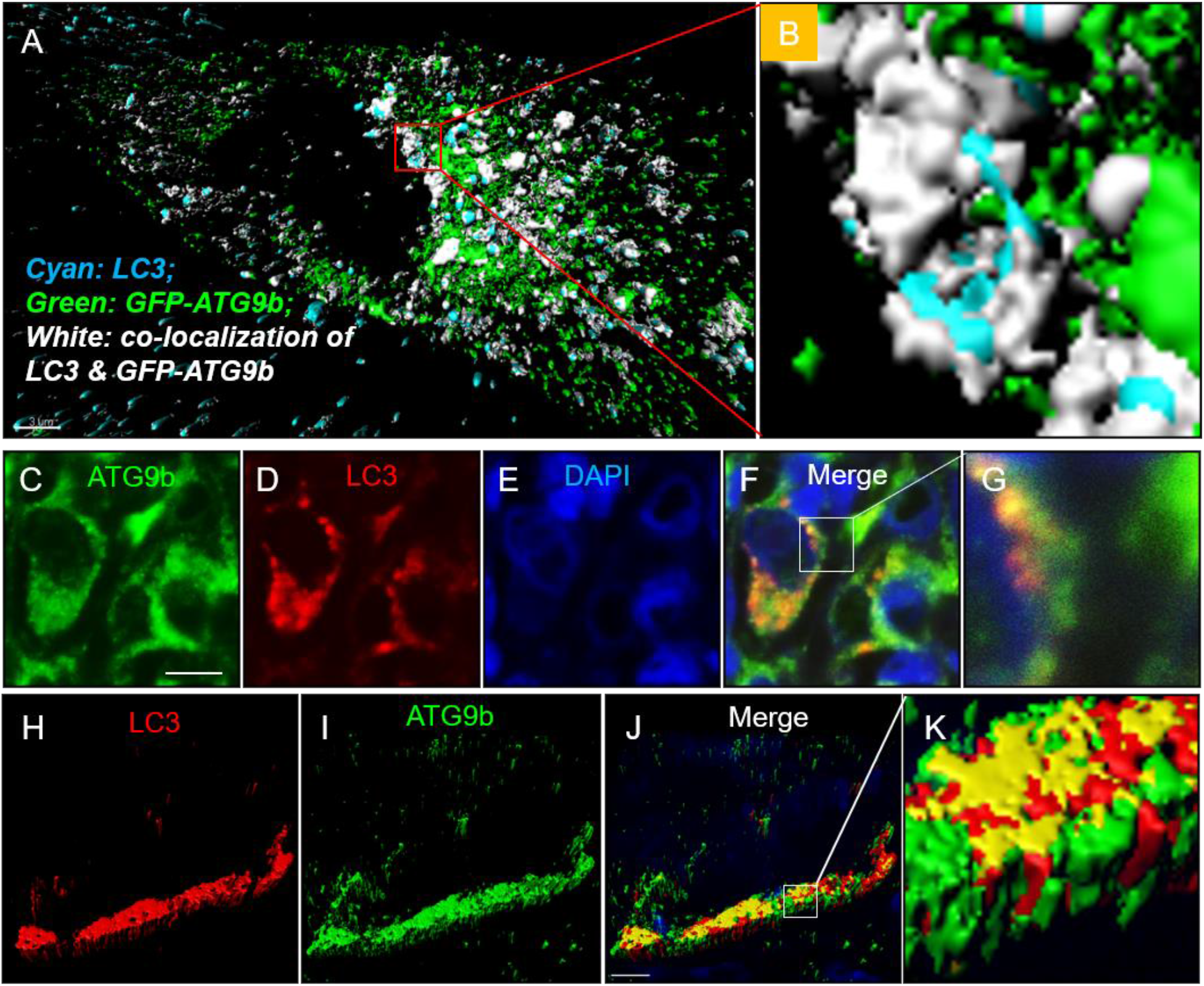
ATG9b co-localizes with LC3. A and B: H1299 cells were transiently transfected with GFP-ATG9b, stained with anti-LC3, and subjected to SIM structure analysis. A shows the regenerated structures that include single channel surfaces for GFP (green) and LC3 (cyan) and the GFP and LC3 co-localized surface (white). B is the blown-up of a red boxed area from A as indicated. Bar: 3 μm. C-G: confocal fluorescence microscopy images for parental HCC827 cells stained with anti-ATG9b (green), anti-LC3 (red), DAPI and (blue). G is the blown-up of the white boxed area from F as indicated. Bar: 5 μm. H-K: regenerated SIM structures for parental HCC827 cells stained with anti-ATG9b (green) and anti-LC3 (red). K is the blown-up of the white boxed area from J as indicated. Bar: 5 μm.

**Supplementary Fig. 2.**
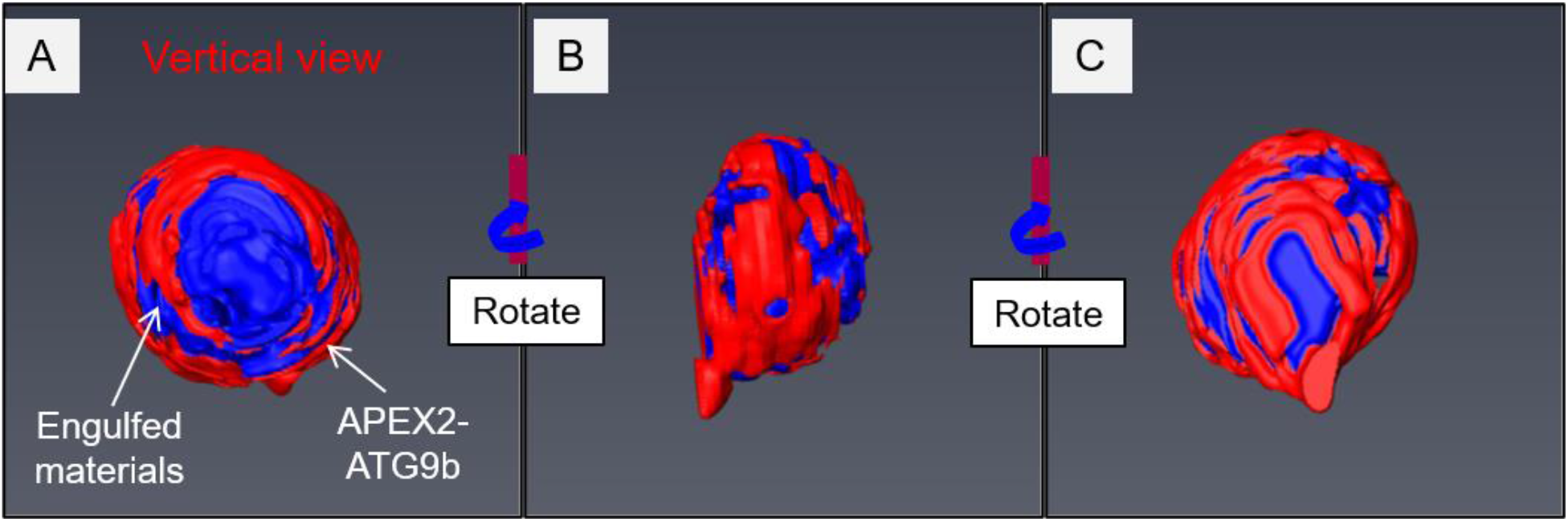
The autophagosome membrane localization of ATG9 in the SBF-SEM structure. H1299 cells were transiently transfected with APEX2-ATG9b and subjected to SBF-SEM analysis. A: the regenerated SBF-SEM autophagosome structure showing the autophagosome membrane localization of the APEX2-ATG9b. B and C: the same structures in A were rotated to provide more comprehensive views.

**Supplementary Fig. 3.**
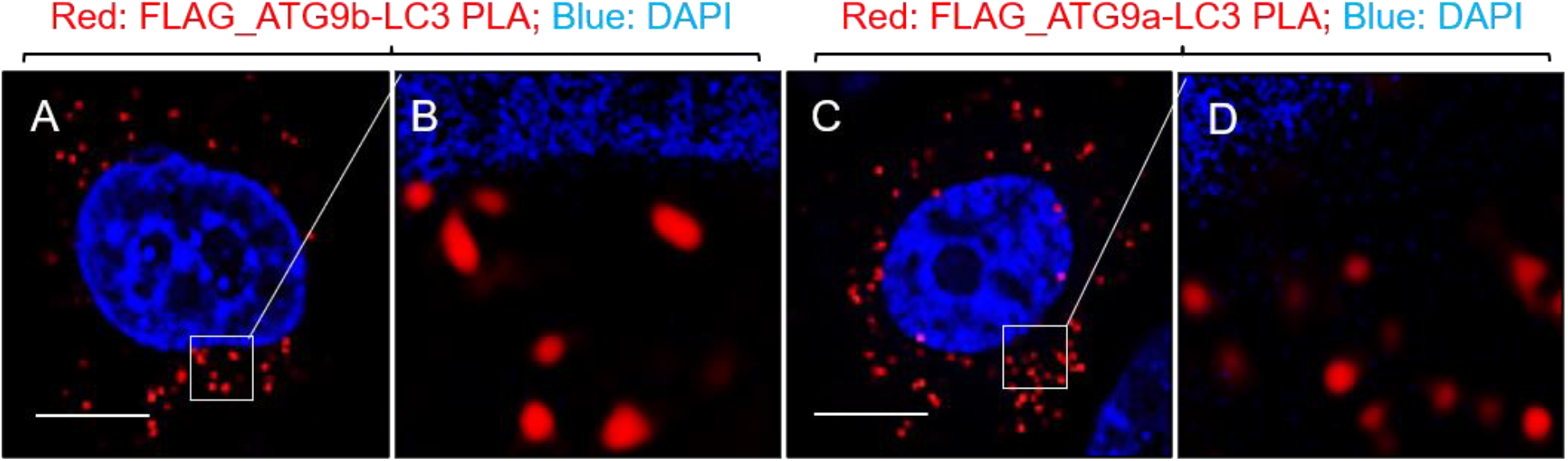
The maATG9s bind to LC3 in situ. H1299 cells were transiently transfected with FLAG-ATG9b (A and B) or FLAG-ATG9a (C and D) and stained with DAPI (blue, to indicate the nucleus. Red dots indicate the sites where maATG9s bind to LC3. B and D are blown-ups of the white boxed areas from A and C, respectively. Bars: 5 μm.

**Supplementary Fig. 4.**
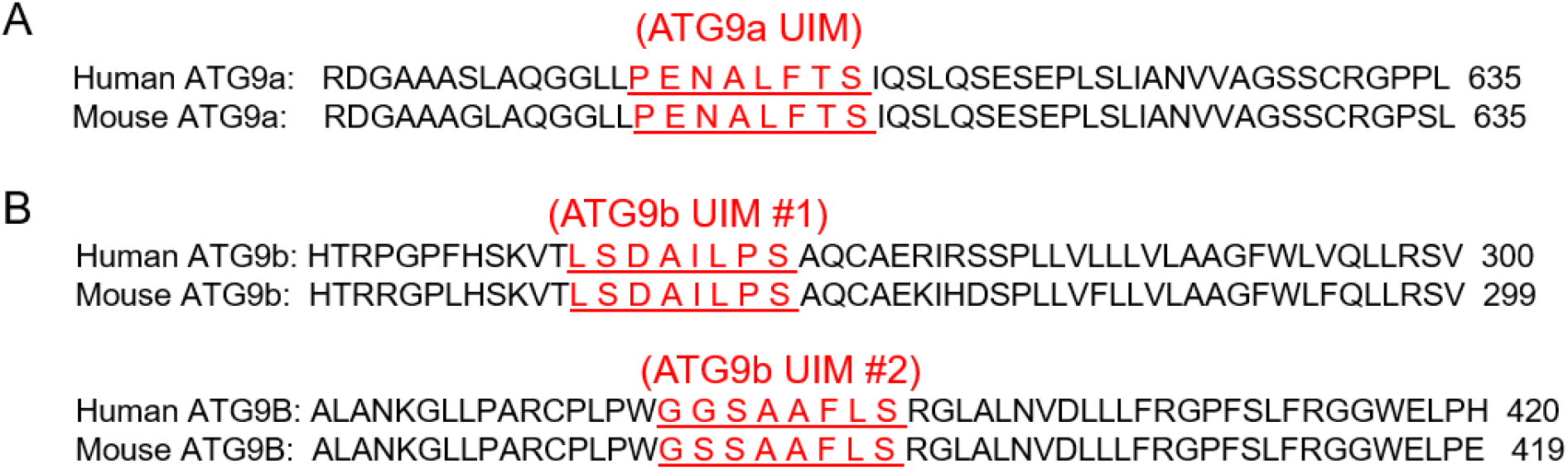
The protein amino acid sequences of ATG9a (A) and ATG9b (B) containing UIM motives.

**Supplementary Fig. 5.**
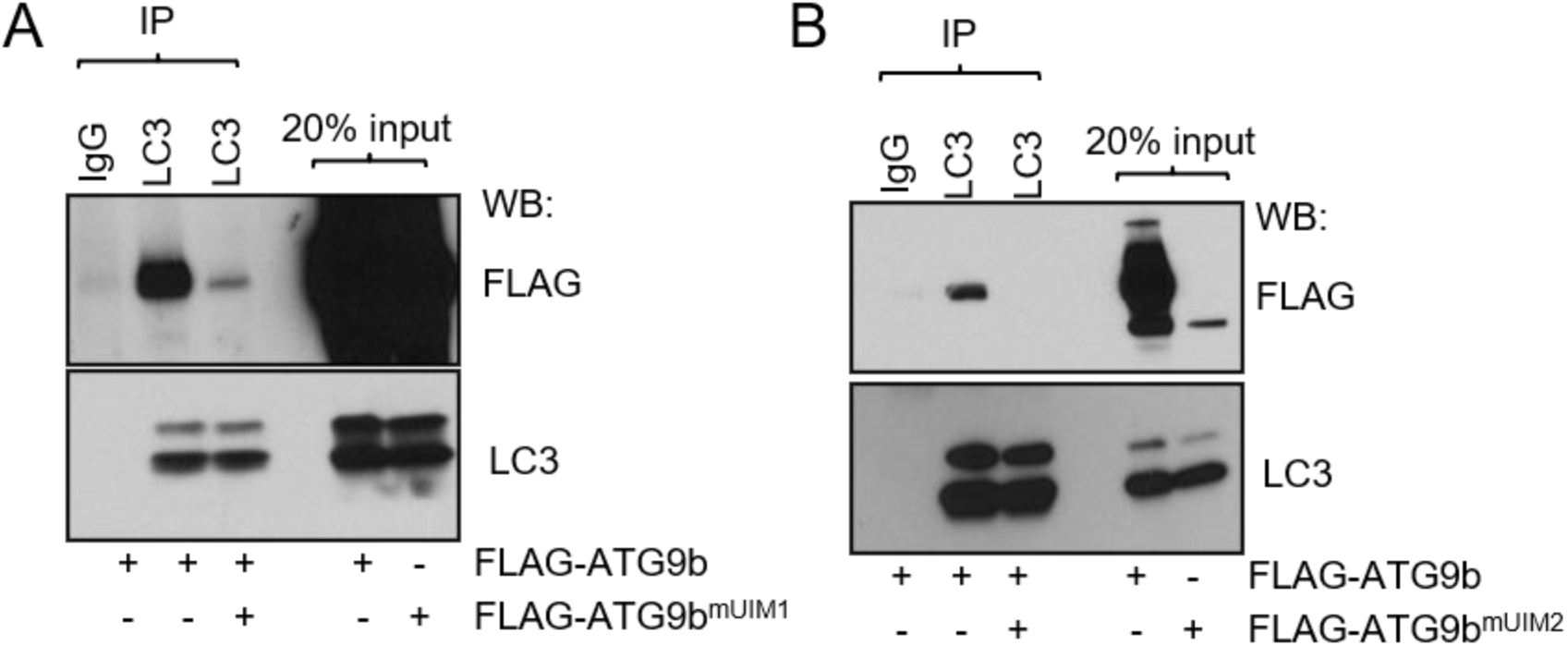
IP-Western blots for H1299 cells transiently co-transfected with FLAG-ATG9b and FLAG-ATG9b^mUIM1^ (A) or FLAG-ATG9b^mUIM1^ (B).

**Supplementary Fig. 6.**
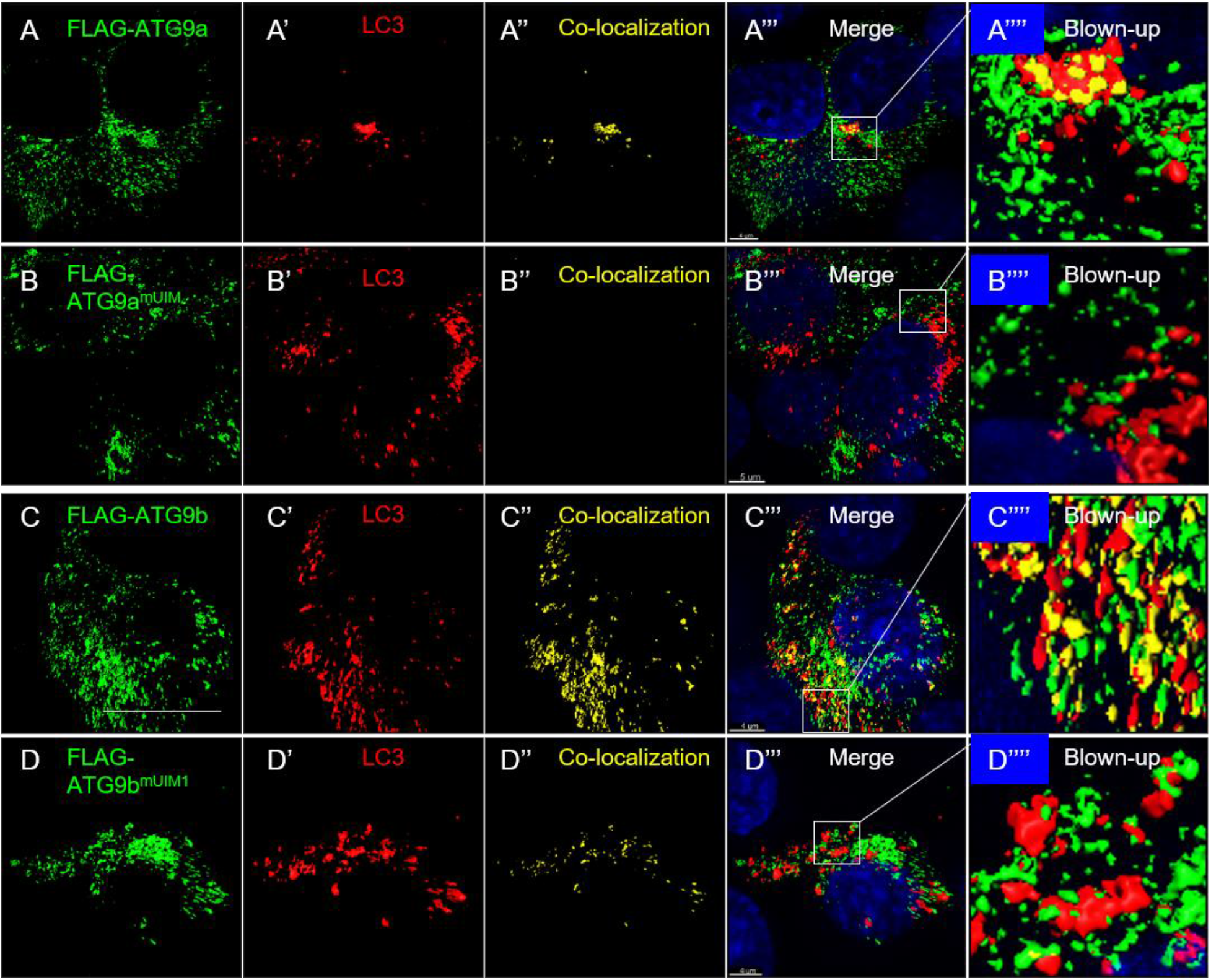
The localization of maATG9s to the autophagosome depends on UIM motives. H1299 human lung cancer cells were transiently transfected with FLAG-ATG9a (A-A’’’’), FLAG-ATG9a^mUIM^ (B-B’’’’), FLAG-ATG9b (C-C’’’’), and FLAG-ATG9b^mUIM1^ (D-D’’’’), stained with FLAG (A, B, C, and D) and LC3 (A’, B’, C’, and D’), and subjected to SIM structure analysis. A, B, C, D, A’, B’, C’, and D’: regenerated single channel surfaces for Flag and LC3 as indicated. A’’, B’’, C’’, and D’’: regenerated surfaces for the co-colocalizations of LC3 and FLAG. A’’’, B’’’, C’’’, and D’’’: the merged SIM structures. DAPI staining was included to show the nuclei. A’’’’, B’’’’, C’’’’, and D’’’’: the blown-up for the white boxed areas from A’’’, B’’’, C’’’, and D’’’ as indicated. Bars: 4 μm in A’’’, C’’’, and D’’’, and 5 μm in B’’’.

**Supplementary Fig. 7.**
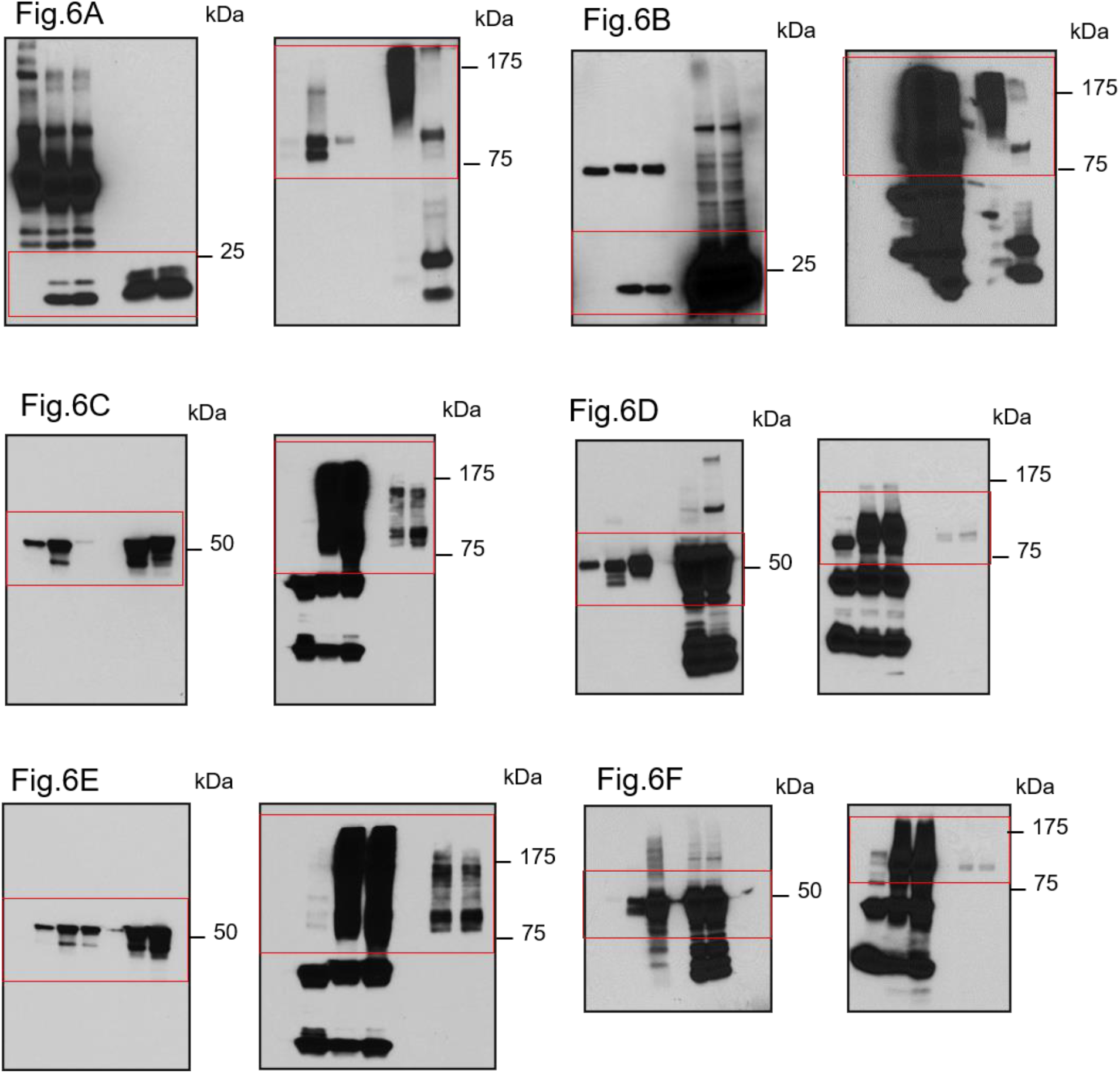
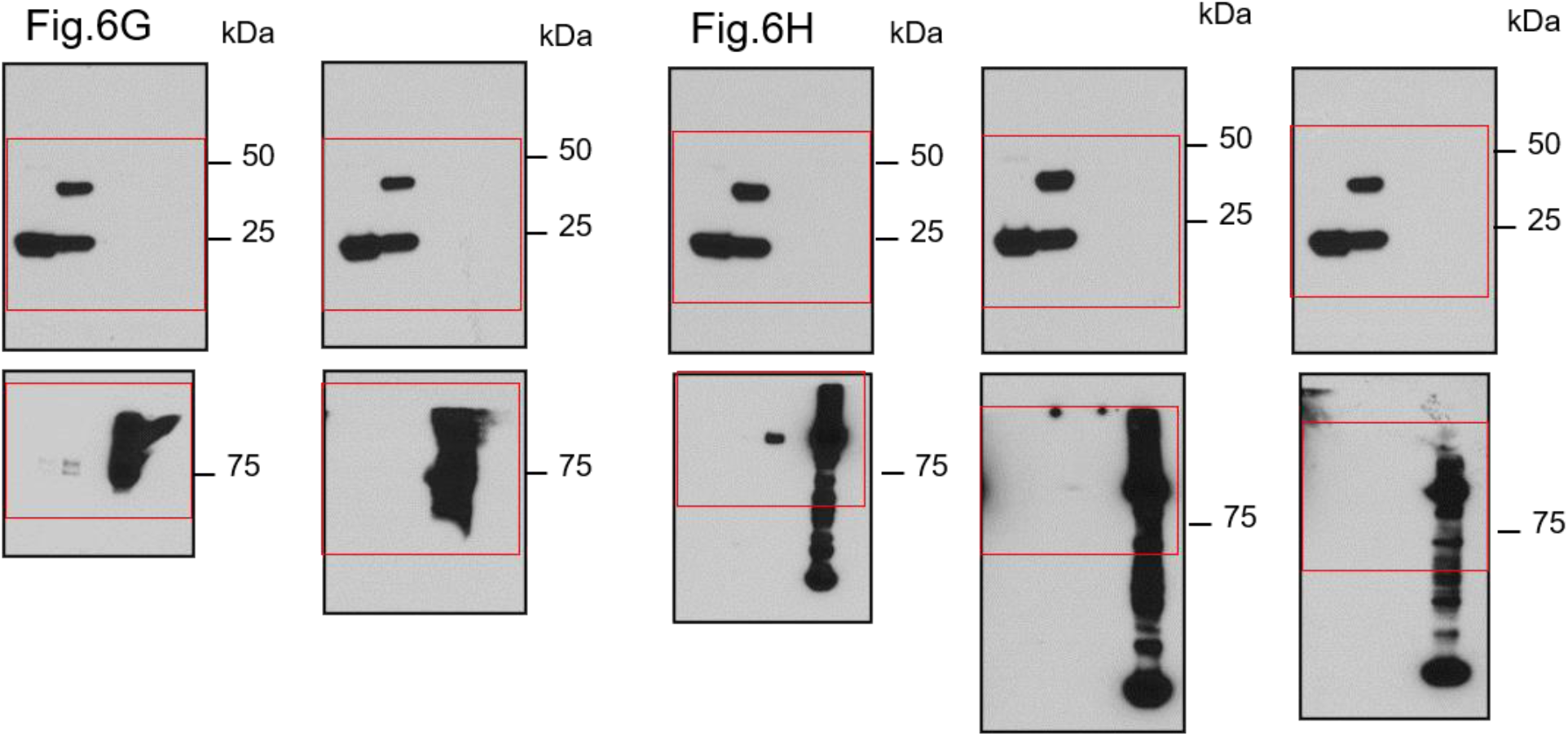
Uncut gels for Fig.6

**Supplementary Fig. 8.**
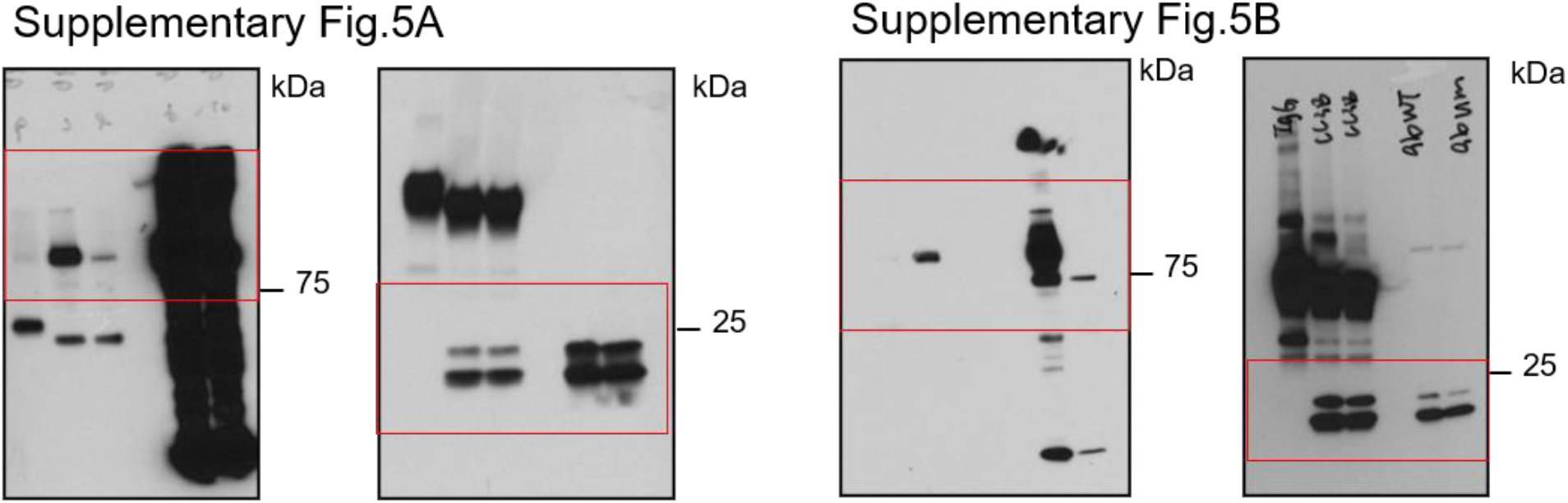
Uncut gels for Supplementary Fig.5

## SUPPLEMENTARY VIDEOS

**Supplementary Video 1.**
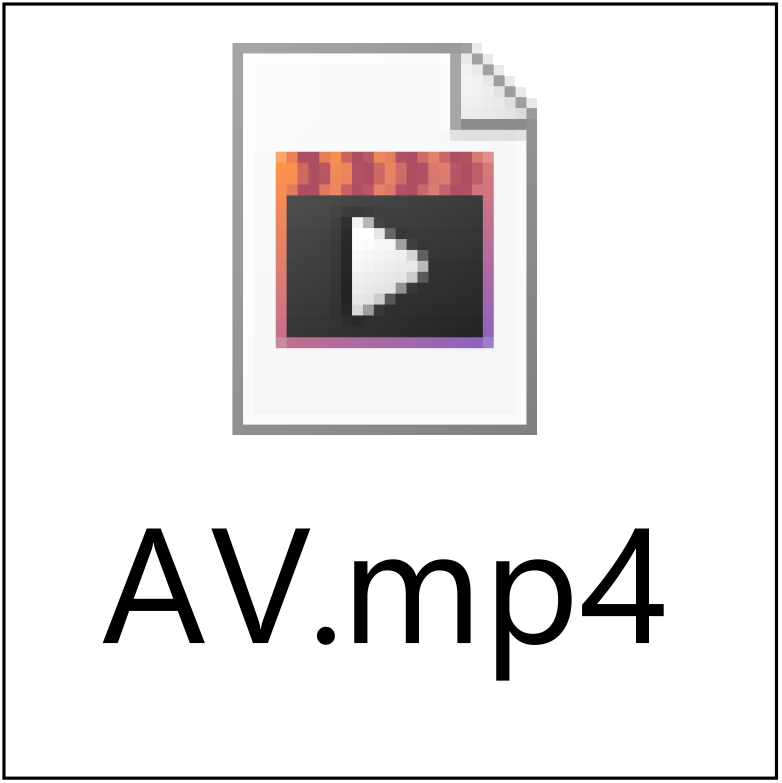
Images shown in Fig.5A-O are used to generate this MP4 video to show a serially sectioned autophagosome.

---

**Supplementary Table 1.**
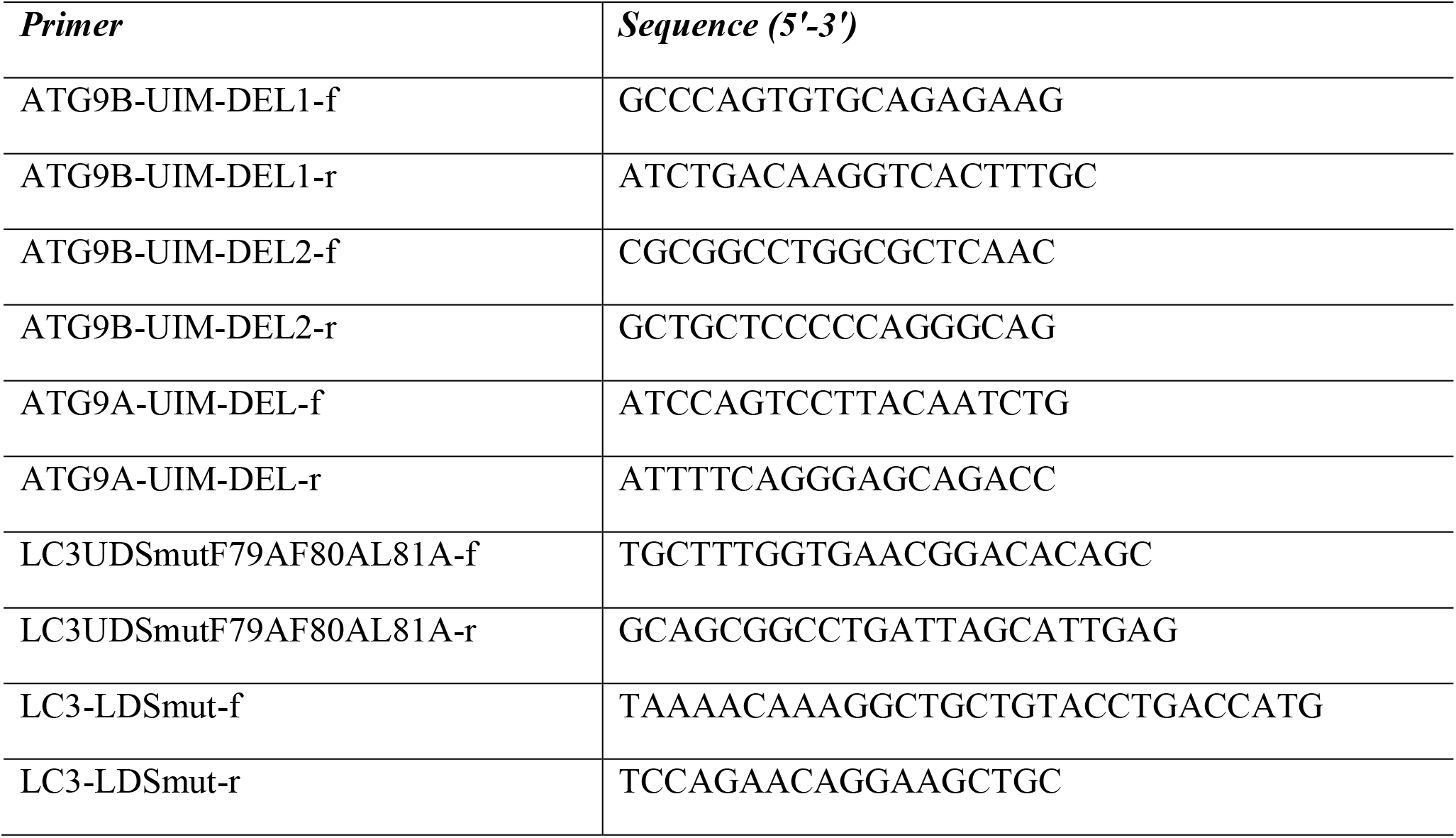
Information for the primers used in this study.

**Supplementary Table 2.** Information for the antibodies used in this study

| *Antibody* | *Source* | *Catalog number* | *Application* |
| --- | --- | --- | --- |
| EEA1 | Cell Signaling | Rabbit mAb 3288 | IF |
| RCAS1 | Cell Signaling | Rabbit mAb 12290 | IF |
| E-cadherin | Cell Signaling | Rabbit mAb 3195 | IF |
| PDI | Cell Signaling | Rabbit mAb 3501 | IF |
| LAMP1 | Cell Signaling | Rabbit mAb 9091 | IF |
| LC3 | Cell Signaling | Rabbit mAb 12741 | IF |
| FLAG | Sigma | F1804 | IF, IP, Western blotting, PLA assay |
| WIPI2 | Abcam | ab105459 | IF |
| LC3 | Cell Signaling | Rabbit mAb 3868 | IP, Western blotting, PLA assay |
| GFP | Santa Cruz | Mouse mAb sc9996 | IP, Western blotting |
| GST | Cell Signaling | Rabbit pAb 2622 | Pull-down assay, Western blotting |
| ATG9a | Cell Signaling | Rabbit mAb 13509 | Western blotting |
| Tubulin | Cell Signaling | Rabbit mAb 2125 | Western blotting |
| ATG9b | Thermo Scientific | PA5-78672 | IF |
| Alexa Fluor® 350 Goat<br>Anti-Mouse IgG (H+L) | Invitrogen | A11045 | IF |
| Alexa Fluor® 594 Goat<br>Anti-Rabbit IgG (H+L) | Invitrogen | A11012 | IF |
| Alexa Fluor® 488 Anti-Mouse IgG (H+L) | Cell Signaling | 4408 | IF |
| anti-Mouse HRP | Cell Signaling | 7076 | Western blotting |
| anti-Rabbit HRP | Cell Signaling | 7074 | Western blotting |
| Normal Mouse IgG | Cell Signaling | 5145 | IP |
| Normal Rabbit IgG | Cell Signaling | 2729 | IP |
| LC3 | Cell Signaling | Mouse mAb 83506 | IF, PLA assay |

## REFERENCES

1. Mizushima N. A brief history of autophagy from cell biology to physiology and disease. Nat Cell Biol. 2018May;20(5):521–527. doi: 10.1038/s41556-018-0092-5. Epub 2018 Apr 23. Review. PMID: 29686264

2. Saha S, Panigrahi DP, Patil S, Bhutia SK. Autophagy in health and disease: A comprehensive review. Biomed Pharmacother. 2018Aug;104:485–495. doi: 10.1016/j.biopha.2018.05.007. Epub 2018 May 25. Review. PMID: 29800913

3. Onorati AV, Dyczynski M, Ojha R, Amaravadi RK. Targeting autophagy in cancer. Cancer. 2018Aug;124(16):3307–3318. doi: 10.1002/cncr.31335. Epub 2018 Apr 19. Review. PMID: 29671878

4. Klionsky DJ et al. Guidelines for the use and interpretation of assays for monitoring autophagy (3rd edition). Autophagy. 2016;12(1):1–222. doi: 10.1080/15548627.2015.1100356. PMID: 26799652 PMCID: PMC4835977

5. Galluzzi L et al. Molecular definitions of autophagy and related processes. EMBO J. 2017 Jul 3; 36(13):1811–1836. doi: 10.15252/embj.201796697. Epub 2017 Jun 8. PMID: 28596378 PMCID: PMC5494474

6. Clarke AJ, Simon AK. Autophagy in the renewal, differentiation and homeostasis of immune cells. Nat Rev Immunol. 2018 Dec 7. doi: 10.1038/s41577-018-0095-2. [Epub ahead of print] Review. PMID: 30531943

7. Lamb CA, Yoshimori T, Tooze SA. The autophagosome: origins unknown, biogenesis complex. Nat Rev Mol Cell Biol. 2013Dec;14(12):759–74. Epub 2013 Nov 8. Review. PMID: 24201109

8. Grasso D, Renna FJ, Vaccaro MI. Initial Steps in Mammalian Autophagosome Biogenesis. Front Cell Dev Biol. 2018 Oct 23;6:146. doi: 10.3389/fcell.2018.00146.eCollection 2018. PMID: 30406104

9. Orsi A, Razi M, Dooley HC, Robinson D, Weston AE, Collinson LM, Tooze SA. Dynamic and transient interactions of Atg9 with autophagosomes, but not membrane integration, are required for autophagy. Mol Biol Cell. 2012May;23(10):1860–73. doi: 10.1091/mbc.E11-09-0746. Epub 2012 Mar 28. PMID: 22456507

10. Ungermann C, Reggiori F. Atg9 proteins, not so different after all. Autophagy. 2018;14(8):1456–1459. doi: 10.1080/15548627.2018.1477382. Epub 2018 Jul 23. PMID: 29966469

11. Noda T. Autophagy in the context of the cellular membrane-trafficking system: the enigma of Atg9 vesicles. Biochem Soc Trans. 2017 Dec 15;45(6):1323–1331. doi: 10.1042/BST20170128. Epub 2017 Nov 17. Review. PMID: 29150528

12. Puri C, Renna M, Bento CF, Moreau K, Rubinsztein DC. Diverse autophagosome membrane sources coalesce in recycling endosomes. Cell. 2013 Sep 12;154(6):1285–99. doi: 10.1016/j.cell.2013.08.044. PMID: 24034251

13. Puri C, Renna M, Bento CF, Moreau K, Rubinsztein DC. ATG16L1 meets ATG9 in recycling endosomes: additional roles for the plasma membrane and endocytosis in autophagosome biogenesis. Autophagy. 2014Jan;10(1):182–4. doi: 10.4161/auto.27174. Epub 2013 Nov 19. PMID: 24257061

14. Mercer TJ, Gubas A, Tooze SA. A molecular perspective of mammalian autophagosome biogenesis. J Biol Chem. 2018 Apr 13;293(15):5386–5395. doi: 10.1074/jbc.R117.810366. Epub 2018 Jan 25. Review. PMID: 29371398

15. Zhuang X, Chung KP, Luo M, Jiang L. Autophagosome Biogenesis and the Endoplasmic Reticulum: A Plant Perspective. Trends Plant Sci. 2018Aug;23(8):677–692. doi: 10.1016/j.tplants.2018.05.002. Epub 2018 Jun 18. Review. PMID: 29929776

16. Webber JL, Tooze SA. Coordinated regulation of autophagy by p38alpha MAPK through mAtg9 and p38IP. EMBO J. 2010 Jan 6;29(1):27–40. doi: 10.1038/emboj.2009.321. Epub 2009 Nov 5. PMID: 19893488

17. Zavodszky E, Vicinanza M, Rubinsztein DC. Biology and trafficking of ATG9 and ATG16L1, two proteins that regulate autophagosome formation. FEBS Lett. 2013 Jun 27;587(13):1988–96. doi: 10.1016/j.febslet.2013.04.025. Epub 2013 May 11. Review. PMID: 23669359

18. Søreng K, Munson MJ, Lamb CA, Bjørndal GT, Pankiv S, Carlsson SR, Tooze SA, Simonsen A. SNX18 regulates ATG9A trafficking from recycling endosomes by recruiting Dynamin-2. EMBO Rep. 2018Apr;19(4). pii: e44837. doi: 10.15252/embr.201744837. Epub 2018 Feb 7. PMID: 29437695

19. Engelsberg A, Hermosilla R, Karsten U, Schülein R, Dörken B, Rehm A. The Golgi protein RCAS1 controls cell surface expression of tumor-associated O-linked glycan antigens. J Biol Chem. 2003 Jun 20;278(25):22998–3007. Epub 2003 Apr 2. PMID: 12672804

20. González-Mariscal L, Miranda J, Gallego-Gutiérrez H, Cano-Cortina M, Amaya E. Relationship between apical junction proteins, gene expression and cancer. Biochim Biophys Acta Biomembr. 2020 Mar 30:183278. doi: 10.1016/j.bbamem.2020.183278. [Epub ahead of print] PMID: 32240623

21. Coelho JP, Feige MJ. In case of stress, hold tight: phosphorylation switches PDI from an oxidoreductase to a holdase, tuning ER proteostasis. EMBO J. 2020 Apr 2:e104880. doi: 10.15252/embj.2020104880. [Epub ahead of print] PMID: 32239769

22. Lam SS, Martell JD, Kamer KJ, Deerinck TJ, Ellisman MH, Mootha VK, Ting AY. Directed evolution of APEX2 for electron microscopy and proximity labeling. Nat Methods. 2015Jan;12(1):51–4. doi: 10.1038/nmeth.3179. Epub 2014 Nov 24. PMID: 25419960

23. Kerstens W, Kremer A, Holtappels M, Borghgraef P, Lippens S, Van Dijc P.. Three-Dimensional Visualization of APEX2-Tagged Erg11 in Saccharomyces cerevisiae Using Focused Ion Beam Scanning Electron Microscopy. mSphere. 2020 Feb 5;5(1). pii: e00981–19. doi: 10.1128/mSphere.00981-19. PMID: 32024705

24. Shi Y, Wang L, Zhang J, Zhai Y, Sun F. Determining the target protein localization in 3D using the combination of FIB-SEM and APEX2. Biophys Rep. 2017;3(4):92–99. doi: 10.1007/s41048-017-0043-x. Epub 2017 Nov 4. PMID: 29238746

25. Kläsener K, Yang J, Reth M. Study B Cell Antigen Receptor Nano-Scale Organization by In Situ Fab Proximity Ligation Assay. Methods Mol Biol. 2018;1707:171–181. doi: 10.1007/978-1-4939-7474-0_12. Review. PMID: 29388107

26. Greenwood C, Johnson G, Dhillon HS, Bustin S. Recent progress in developing proximity ligation assays for pathogen detection. Expert Rev Mol Diagn. 2015;15(7):861–7. doi: 10.1586/14737159.2015.1044440. Epub 2015 May 8. Review. PMID: 25955213

27. Marshall RS, Hua Z, Mali S, McLoughlin F, Vierstra RD. ATG8-Binding UIM Proteins Define a New Class of Autophagy Adaptors and Receptors. Cell. 2019 Apr 18;177(3):766–781.e24. doi: 10.1016/j.cell.2019.02.009. Epub 2019 Apr 4. PMID: 30955882

28. Yamamoto H, Kakuta S, Watanabe TM, Kitamura A, Sekito T, Kondo-Kakuta C, Ichikawa R, Kinjo M, Ohsumi Y. Atg9 vesicles are an important membrane source during early steps of autophagosome formation. J Cell Biol. 2012 Jul 23;198(2):219–33. doi: 10.1083/jcb.201202061. PMID: 22826123

29. Lee IH, Cao L, Mostoslavsky R, Lombard DB, Liu J, Bruns NE, Tsokos M, Alt FW, Finkel T. A role for the NAD-dependent deacetylase Sirt1 in the regulation of autophagy. Proc Natl Acad Sci U S A. 2008 Mar 4. 105(9):3374–9. 10.1073/pnas.0712145105 PubMed 18296641

30. Koyama-Honda I, Itakura E, Fujiwara TK, Mizushima N. Temporal analysis of recruitment of mammalian ATG proteins to the autophagosome formation site. Autophagy. 2013Oct;9(10):1491–9. doi: 10.4161/auto.25529. Epub 2013 Jul 10.10.4161/auto.25529 PubMed 23884233

